# Lower dormancy with rapid germination is an important strategy for seeds in an arid zone with unpredictable rainfall

**DOI:** 10.1101/660159

**Authors:** Corrine Duncan, Nick Schultz, Wolfgang Lewandrowski, Megan Good, Simon Cook

## Abstract

Seed germination traits are key drivers of population dynamics, yet they are under-represented in community ecology studies, which have predominately focussed on adult plant and seed morphological traits. We studied the seed traits and germination strategy of eight woody plant species to investigate regeneration strategies in the arid zone of eastern Australia. To cope with stochastic and minimal rainfall, we predict that arid seeds will either have rapid germination across a wide range of temperatures, improved germination under cooler temperatures, or dormancy and/or longevity traits to delay or stagger germination across time. To understand how temperature affects germination responses, seeds of eight keystone arid species were germinated under laboratory conditions, and under three diurnal temperatures (30/20°C, 25/15°C and 17/7°C) for 30 days. Seeds of species in this study are currently stored for minesite restoration projects, hence we tested for decline in seed viability across 24 months in dry storage at similar storage conditions (≈20°C). Six of the eight arid species studied had non-dormant, rapidly germinating seeds, and only two species had physiological dormancy traits. Seed longevity differed widely between species, from one recalcitrant species surviving only months in storage (P50 = <3 months) and one serotinous species surviving for many years (P50 = 84 months). Our results highlight the importance of understanding the reproductive strategies of plant species in arid environments. Rapid germination, the dominant seed trait of species included in this study, allows arid species to capitalise on sporadic rainfall. However, some species also exhibit dormancy and delayed germination; this an alternative strategy which spreads the risk of germination failure over time. We highlight important seed traits and germination strategies of plants from an arid zone with stochastic rainfall and discuss the implications for their restoration.

## Introduction

Seed traits and germination strategies drive plant community dynamics and provide insight into species’ adaptations to environmental filters [1] and community composition [2]. Despite this, seed traits are under-represented in community ecology studies [3-5]. Knowledge of seed traits and germination strategies is necessary to describe plant niches, to anticipate population dynamics under changes in land use [6], and to assess plant responses to the environment [7]. By studying seed traits and germination responses we can obtain ecologically meaningful data about the functional properties of plant communities that improve predictions of plant assemblages under natural, and anthropogenic, environmental change [8].

Seed traits and germination strategies, which are often unrelated to other plant traits [9], can inform us about the reproductive strategies of species occurring in particular environments [10]. Seed mass is the most widely studied seed trait, and there are strong correlations among seed mass, germination rate, survival and establishment [11, 12]. Small seeds generally germinate faster than heavy seeds [13], however the chance of survival and establishment at the seedling or later plant stages is greater for heavier seeds [14, 15]. Furthermore, small-seeded species are generally able to produce more seeds [16]. Physiological responses of seeds to environmental cues enable germination to occur when conditions are most suitable for seedling establishment [17, 18]. Conversely, seed death can occur if germination occurs in unfavourable environmental conditions; thus germination timing has strong fitness consequences [19-21].

The primary abiotic filters for recruitment of plant communities are water availability and the season it occurs in, and arid seeds exhibit adaptive traits to tolerate or avoid drought conditions [22, 23]. Rainfall events that last several days are rare in arid zones, and smaller rainfall events are likely to result in the drying of upper soil layers before the germination process is complete [24], causing seedling mortality after germination. Conversely, a rainfall event during winter may present slower evaporation rates from the soil surface than during summer. Hence, for species that rely on avoidance strategies to survive drought conditions, the timing of recruitment must coincide with periods of reliable soil water availability [25]. Dormancy is an important drought avoidance mechanism for arid species as it restricts germination during unsuitable environmental conditions [26], whereas rapid germination is a drought tolerance strategy to facilitate the rapid exploitation of temporarily favourable conditions [27]. The evolution towards mature seeds with less nutritive tissue available to the embryo has enabled seeds to germinate faster [28, 29] and emerge when rainfall events are short.

The level of seed dormancy can vary among seeds within a population so that germination of individuals occurs over several seasons [30-32]. This reproductive strategy has been termed bet-hedging, as it limits synchronous germination events, spreading the risk of germination failure across seasons. This increases long-term fitness by ensuring the entire seed cohort are not lost during unfavourable conditions [33] provided seeds survive through multiple seasons of unfavourable conditions. Hence, dormancy and longevity are key traits of bet-hedging. However, in unpredictable environments, selection may favour plants that can reproduce rapidly and frequently, and hence bet-hedging may be less prevalent.

In arid ecosystems, Baskin and Baskin (34) estimate that 85% of plant species produce dormant seeds. The prevalence of dormancy generally increases with aridity [35], but the influence of rainfall predictability on seed traits is not as well studied. Modelling by Brown and Venable (36) predicts that the prevalence of dormancy traits should increase with decreasing rainfall predictability. However, Harel, Holzapfel (37) found that dormancy decreased with rainfall predictability in desert annuals. Arid seeds are often characterised by faster germination rates than those from temperate ecosystems [38, 39] after dormancy is overcome. Species that germinate quickly are able to utilise the short pulses of water availability, reducing the likelihood of seed mortality [40], while the seedlings of slower-germinating species may be limited to using dwindling water availability at the end of longer rainfall pulses [41]. Further empirical evidence is required from a greater suite of species (particularly perennials), and from a greater range of environments, to test the effect of rainfall predictability on seed traits and to determine the prevalence of dormancy in the arid zone.

This study investigates the seed traits and germination strategy of eight keystone, Australian arid zone species. We focused on traits that may be critical to species’ persistence in arid ecosystems, and that are essential to understand if we are to successfully restore these species using seed. We hypothesise that seeds will be categorised by one of three strategies beneficial to recruitment in arid zones. To cope with stochastic and low rainfall, we predict that arid seeds will either 1) have low seed mass but with rapid germination, across a wide range of temperatures, to ensure seedling establishment before soil moisture evaporates, 2) show improved germination under cool, winter temperatures where soil evaporation rates are lowest, or 3) have dormancy and/or longevity traits to delay or stagger germination, therefore spreading the risk of germination failure across time. Specifically, we measure seed dormancy and embryo traits, germination responses under different temperature regimes, and seed longevity under ambient storage conditions. We also test if seed mass, and other traits, are related to germination strategy and whether seed traits can be used as a proxy for germination strategy. We highlight important germination strategies of plants from an arid zone with stochastic rainfall, and discuss seed traits that may impact on the success of restoration efforts for these species.

## Materials and methods

### Study species and seed collection

We chose the following Australian arid zone species for this study: trees *Casuarina pauper* F.Muell. ex L.A.S.Johnson (Casuarinaceae), *Myoporum platycarpum* ssp. *platycarpum* R.Br., *Geijera parviflora* Lindl. (Scrophulariaceae), *Alectryon oleifolius* ssp. *canescens* S.T.Reynolds (Sapindaceae) and *Hakea tephrosperma* R.Br. (Proteaceae), and understory shrubs from the Chenopodiaceae family, *Atriplex rhagodioides* F.Muell., *Maireana sedifolia* (F.Muell.) Paul G.Wilson and *Maireana pyramidata* (Benth.) Paul G.Wilson. All species in this study are targeted for restoration after mineral sand mining. Vegetation communities prior to clearing include three major plant community types; shrublands dominated by *M. sedifolia*, woodlands dominated by *C. pauper* and *M. platycarpum* and woodlands with shrubland understory, dominated by *C. pauper* and understory shrubs *M. sedifolia* and *M. pyramidata* [42]. Less dominant tree species occur as small, scattered patches across the landscape and include *H. tephrosperma, A. oleifolius* and *G. parviflora*.

The climate of the study area is semi-arid (250 mm mean annual rainfall) although average annual rainfall can often fall below 200 mm for consecutive years. Temperatures range from 2°C to 47°C, with cooler mean daily temperatures from May to August [Fig. 1; 43]. Across 60 years of climate data, average monthly rainfall was 24 mm and, unlike most arid zones across the globe, there is no distinct wet season [43].

**Fig. 1.**
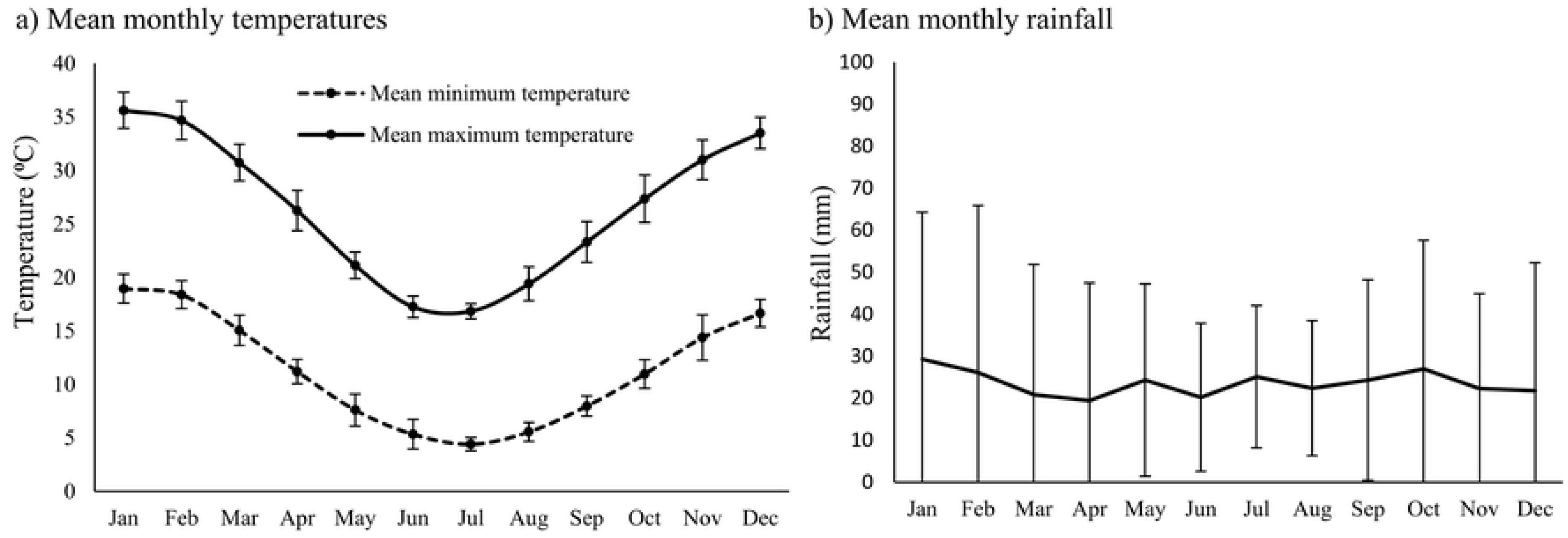
(a) Mean monthly minimum and maximum temperatures from 2003-2016, and (b) mean monthly rainfall, from 1956-2016, at the study site. Unlike most other arid zones, there is no predictable wet season.

Mature seeds of each species were collected from remnant populations within a 200-km radius of a mine site (33°22’05”S, 142°13’36”E) at which restoration of these species is planned. Seeds were either collected by hand, donated from Ecotypic Pty Ltd and Cristal Mining, or purchased from Ogyris Pty Ltd. Where possible, seeds were collected within the six months prior to testing. However, due to lack of seeding events, it was necessary to use seeds stored for over one year in some species (seed ages shown in Table 1). Seeds were manually cleaned and stored in paper bags, under cool dark conditions. Bracts and seed covering structures were removed prior to seed weight measurements and germination experiments.

**Table 1.** Measured seed traits of species in this study. Age = age of seed used in tests; Dry mass = mean weight of 100 seeds; SF = initial seed fill; V = seed viability; ET = embryo type, classified according to Martin (1946); E:S = embryo to shoot ratio (seeds without endosperm indicated as ∼1 and seeds with embryo longer than seed indicated as >1); I72 = seed mass increase at 72hr; and DT = dormancy type classified according to Baskin and Baskin (2014; ND = non dormant, PD = physiological dormancy). The standard errors of means are shown in parentheses.

| Species | Lifeform | Age<br>(months) | Dry mass (g) | SF (%) | V (%) | ET | E:S | I72 (%) | DT |
| --- | --- | --- | --- | --- | --- | --- | --- | --- | --- |
| *Atriplex rhagodioides* | Shrub | 12 | 0.06 (0.001) | 94 (0.6) | 67 (1.1) | Peripheral | >1 | 21 (1.4) | PD |
| *Maireana sedifolia* | Shrub | 2 | 0.13 (0.001) | 97 (0.3) | 93 (0.5) | Peripheral | >1 | 30 (2.5) | ND |
| *Maireana pyramidata* | Shrub | 8 | 0.24 (0.005) | 89 (0.9) | 62 (0.9) | Peripheral | >1 | 16 (0.4) | ND |
| *Casuarina pauper* | Tree | 5 | 0.32 (0.007) | 72 (0.9) | 35 (0.6) | Investing | ~1 | 10 (0.6) | ND |
| *Hakea tephrosperma* | Tree | 16 | 3.31 (0.058) | 96 (0.8) | 96 (0.6) | Investing | ~1 | 91 (1.8) | ND |
| *Alectryon oleifolius* | Tree | 15 | 2.75 (0.111) | 68 (0.8) | 48 (1.2) | Bent | ~1 | 87 (3.4) | ND |
| *Geijera parviflora* | Tree | 2 | 2.95 (0.136) | 94 (0.5) | 88 (1.1) | Spatulate | 0.89 | 29 (0.7) | PD |
| *Myoporum platycarpum* | Tree | 0.5 | 0.26 (0.005) | 20 (0.7) | 13 (0.6) | Straight | ~1 | N/A | ND |

### Seed traits: embryo type, mass, viability and imbibition

For each species, the ratio of embryo length to seed length (E:S) was calculated based on longitudinal dissections and measurements of 50 fully-imbibed seeds [44]. Seed viability was assessed in a 1% solution of 2,3,5-triphenyl tetrazolium chloride (TZ), except for *H. tephrosperma*, for which seeds were incubated on moist filter paper at 30/20°C due to consistently poor TZ stain results; they gave a germination response of 100%. Embryos that only partially absorbed stain were scored as weakly viable (Fig. 2a-c) and were classified as non-viable seeds. To ensure accurate TZ interpretations, results were frequently compared to tests for viability by germinating seeds on filter paper at diurnal temperatures at 30/20°C. Seed mass was determined using the mean of three replicates of 100 seeds, with results then divided by 100 to represent weight (g) per seed.

**Fig. 2.**
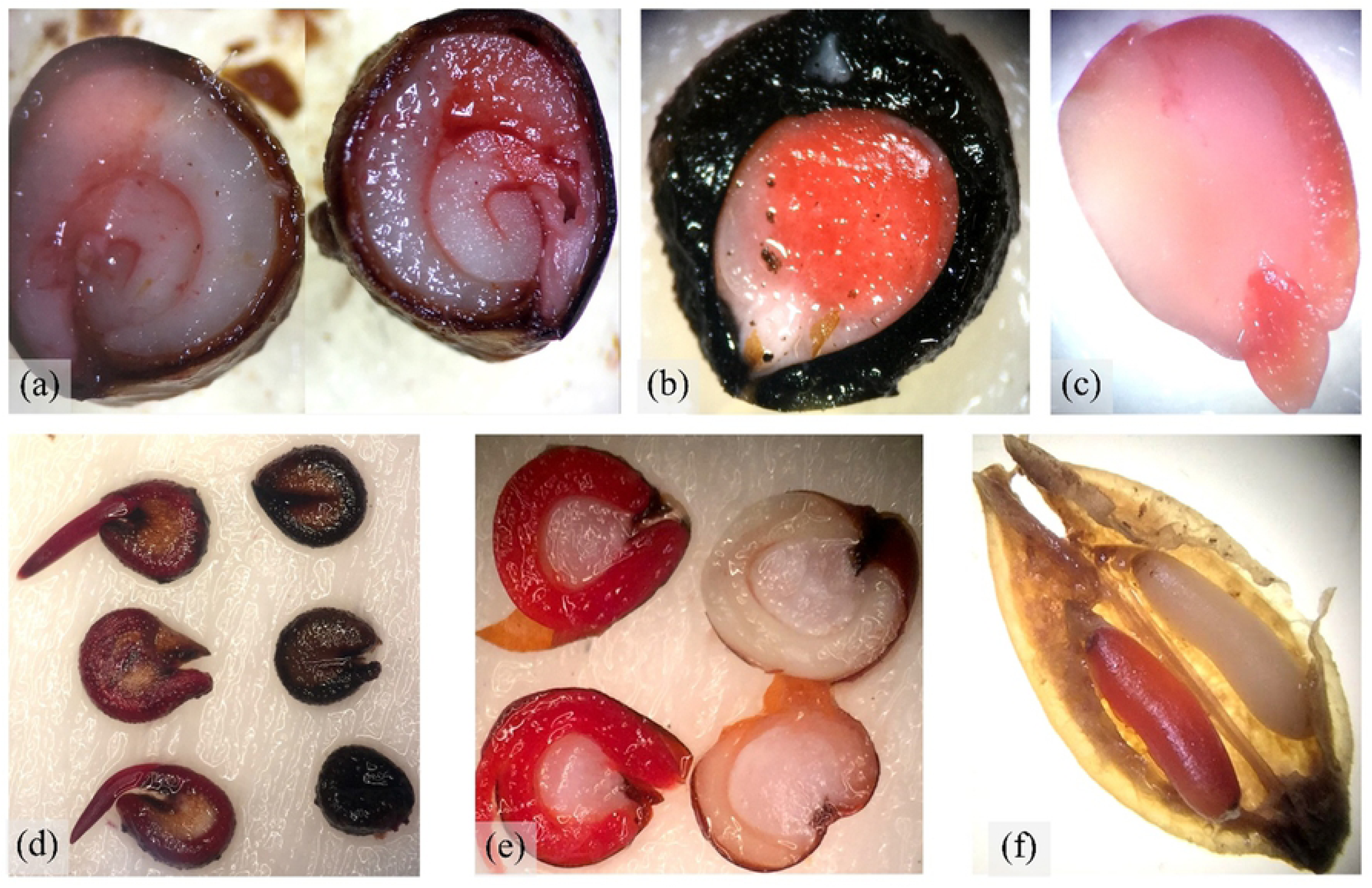
Microscope images of TZ stained seeds for viability tests, including (a) *A. oleifolius*, (b) *G. parviflora*, (c) *C. pauper*, (d) *M. pyramidata*, e) *A. rhagodioides*, and (f) *M. platycarpum*. Weakly viable seeds were only partially stained and considered not viable (a-c). Seeds with viable embryos stained red (left in images d-f) and unviable embryos remained unstained (right-hand-side in d-f).

For imbibition tests, four replicates of 25 seeds were weighed, placed on moist filter paper, and incubated under three diurnal temperatures of 30/20°C, 25/15°C and 17/7°C, under a 12 hr light/dark schedule. Seeds were weighed when dry at beginning of experiment and, after 5 min on moist filter paper, seeds were removed and patted dry with a paper towel to absorb surface moisture, and weighed. To determine increases in seed weight, seeds were re-weighed at 10 min, 30 min and at 1, 2, 3, 6, 9, 24, 48, 72, 96, 120, 144, 168 and 192+ hrs. All bracts and seed coverings were removed for imbibition, germination and viability tests, but they were retained for the longevity experiment as this is how seeds are currently stored after collection. For one species (*G. parviflora*), additional tests were performed in an attempt to alleviate dormancy, including the move-along method (as described by Baskin & Baskin, 2014) and soaking in boiling water and 90% H_2_SO_4_, both treatments for 1 min, 30 min and at 1, 4, 12, 24 and 48 hours.

### Germination responses under diurnal temperatures

Prior to germination treatments, seeds of all species were surface sterilised by soaking in 1% sodium hypochlorite for one minute, then rinsed for 40 seconds with double distilled water. For each species, four replicates of 25 seeds each were used. Seeds were placed in 90-mm diameter petri dishes on filter paper moistened with distilled water and incubated at a 12/12-hr light/dark regime at daily alternating temperatures of 30/20°C, 25/15°C and 17/7°C). Seeds were incubated in cabinets (Thermoline Scientific, temperature and humidity cabinet, Model: TRISLH-495-1-s.d., Vol. 240, Sydney, Australia) under cool-white fluorescent lamps with a 40 μmol.m^-2^ photosynthetic photon flux. To determine the effects of GA_3_ on seed germination, species were incubated at 30/20°C and the filter paper was moistened with a 350 ppm GA_3_ solution. To prevent microbial outbreak and ensure constant hydration during germination tests, seeds were transferred to sterilised Petri dishes weekly, on new filter papers moistened with the same appropriate water/PEG solutions. Seed germination (when the radicle emerged to at least half of seed size) was recorded daily for 30 days, or until germination ceased for four consecutive readings across all treatments. For two species that exhibited dormancy traits (*G. parviflora* and *A. rhagodioides*) an after-ripening treatment was applied by storing seeds for one year, under constant dark and air-conditioned temperatures between 10–20°C and 45–50% humidity. An additional four replicate plates at each diurnal temperature were included for both of these species using the after-ripened seeds. Seeds for the treatments with and without after-ripening were from the same seed lot.

### Longevity

Seeds of species in this study are currently collected for restoration purposes and stored at the study location in air-conditioned shipping containers at ≈20°C, hence we tested the effects of these storage conditions on seed longevity. Seeds were manually cleaned to remove excess organic matter from the seed batch, and stored in paper bags, under constant dark and air-conditioned temperatures between 10–20°C and 45–50% humidity. Temperature and humidity of storage conditions were monitored and recorded once a fortnight for 24 months. To test for viability loss over time in storage, 100 seeds were extracted at 0, 3, 6, 12, 18 and 24 months, and TZ stained as per viability testing methods listed above. Due to seed shortages in *A. oleifolius*, only 60 seeds were tested for viability at each longevity test.

### Data analysis

Imbibition was calculated using increase in seed weight after 72 hours:

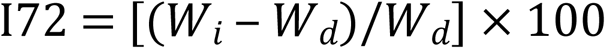

where *W*_*i*_ is mean mass of imbibed seeds and *W*_*d*_ is mean mass of dry seeds [45]. Viability of seed batches was tested in the days prior to experiment, and Viability-Adjusted Germination (VAG) was calculated using the following equation [46]: 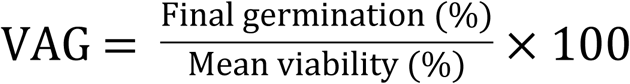

For each replicate dish, the time to minimum germination (T_min_) was taken as the first day that germination was observed, time to 50% germination (T_50_) was the first day that VAG was recorded at ≥50%, and time to maximum germination (T_max_) was the first day at which the maximum VAG was recorded. Mean T_min_, T_50_ and T_max_ were calculated from the four replicates of each species at each diurnal temperature.

Loss of seed viability over time in storage was calculated as:

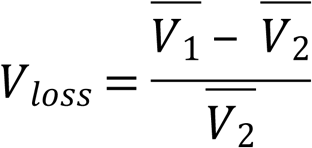

where 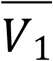 is mean viability at day 0, and 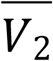 is mean viability at day of test. A Pearson test was used to test the correlation between seed weight and T_50_, P50 and seed fill. One-way ANOVAs were used to test the effects of diurnal temperatures on maximum VAG and T_max_. Shapiro-Wilk tests confirmed the normality of both maximum VAG and T_50_ data prior to ANOVAs. Tukey HSD post-hoc tests were used to make multiple comparsions of means among diurnal temperature treatments. Welsh’s t-tests were performed to compare germination between water and GA_3_ treatents, and to test the effects of after-ripeing in *A. rhagadioides*.

A generalised linear model with binomial error and a probit link function was fitted to the seed longevity data (i.e. loss of viability over time), and thus fit the viability equation [47];

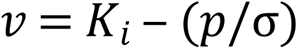

where *v* was the viability after *p* months in storage, σ is the standard deviation of the normal distribution of seed deaths in time, and *K*_*i*_ is the initial seed viability. An estimate of the time taken for seed viability to fall to 50% (P50) was calculated by solving for *p* when *v* = 50%. The Pearson test, Shapiro-Wilk tests, ANOVAs and GLMs were all conducted in R [48].

## Results

There was a weakly significant correlation between seed mass and T_50_ (Fig. 3; P=0.0.034). The two species with the larger seeds exhibited slower germination than the smaller-seeded species. However, there was no correlation between seed mass and P50 (R=0.44, P=0.28) or between seed mass and seed fill (R=0.23, P=0.59).

**Fig. 3.**
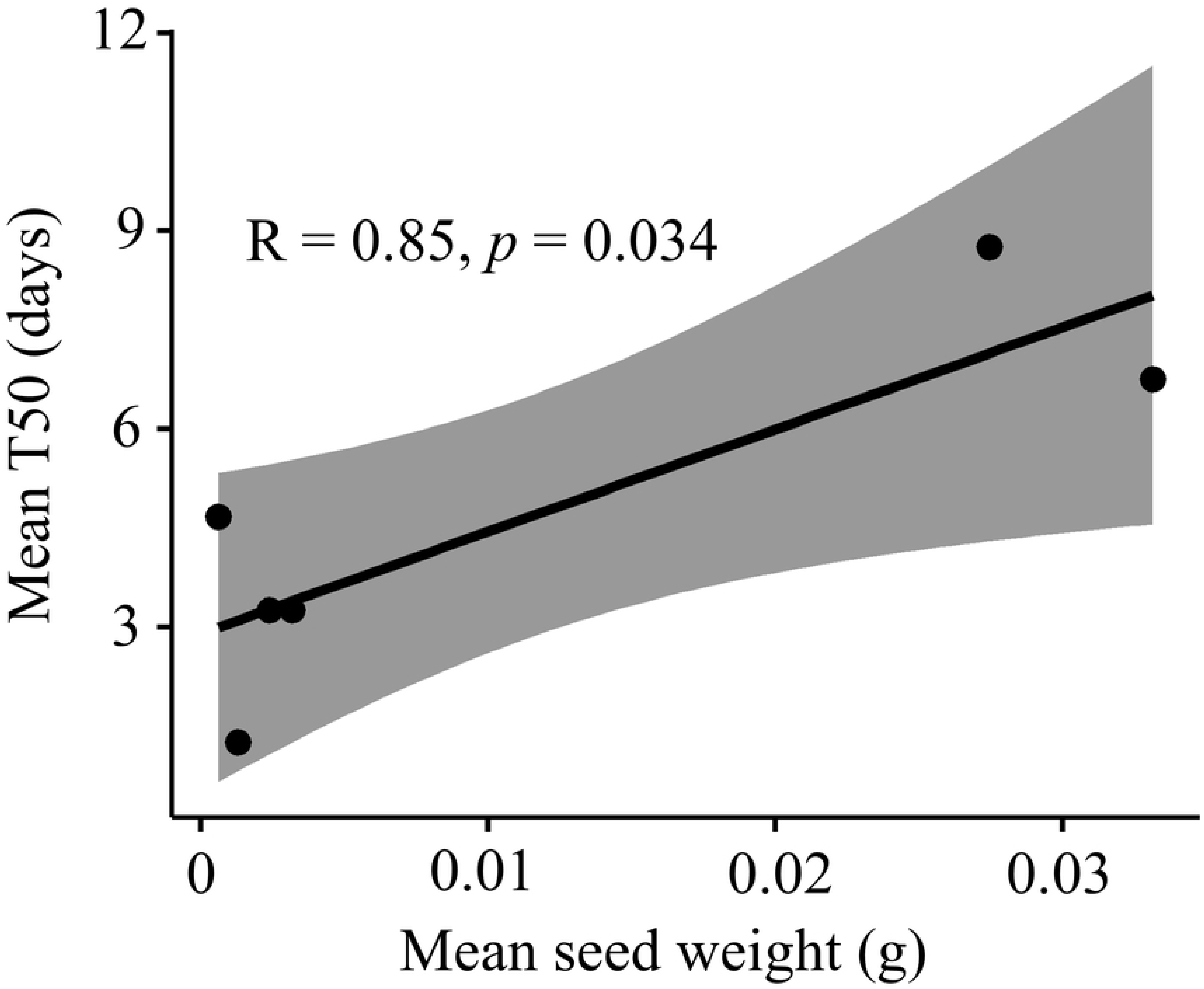
Pearson correlation between mean seed weight and mean T50.

### Viability and dormancy

Each species in the study showed high VAG in at least one diurnal temperature treatment within two weeks and without pre-treatment, except for *A. rhagodioides* and *G. parviflora*. No germination was observed for *G. parviflora* at any of the diurnal temperatures tested, nor through further treatments to relieve dormancy (boiling water and H_2_SO_4_ soaks, GA_3_, after-ripening and the move-along method). Hence the dormancy characteristics of *G. parviflora* were not determined, and no further germination results for this species will be presented. Seed fill was high for all species, except *M. platycarpum*, which was also excluded from germination studies due to lack of viable seeds. Imbibition studies demonstrated that seeds of all species readily take up water within 72 hours of wetting (Table 1).

### Effect of temperature on seed germination proportions

Temperature had no significant effect on maximum VAG for *C. pauper, M. sedifolia* and *A. oleifolius* (P ≥0.2 across all temperatures). However, maximum VAG was significantly lower at the warmest diurnal temperature of 30/20°C for *M. pyramidata* (P = 0.01) and to a lesser degree for *H. tephrosperma* (P = 0.045) and *A. rhagodioides* (P = 0.047; Table 2).

**Table 2.** Mean values for maximum VAG for each species under three diurnal temperatures. Maximum VAG is also shown for the GA_3_ treatment (when diurnal temperature is 30/20°C), and after the after-ripening treatment for *A. rhagodioides*. Letters indicate the results of Tukey pairwise comparisons among the three diurnal temperature treatments. Treatments that share a letter are not significantly different from each other. For the GA_3_ and after-ripening treatments, asterisks represent significant differences compared to the control treatment. For the ‘GA_3_ + after-ripening’ treatment, the comparison is to the GA_3_ only treatment (n.s. = not significant; * 0.05 > p > 0.01; ** 0.01 > p > 0.001; p < 0.001).

| Treatments | *Atriplex*<br>*rhagodioides* | *Maireana*<br>*sedifolia* | *Maireana*<br>*pyramidata* | *Casuarina*<br>*pauper* | *Hakea*<br>*tephrosperma* | *Alectryon*<br>*oleifolius* |
| --- | --- | --- | --- | --- | --- | --- |
| *Diurnal temp* |  |  |  |  |  |  |
| 30/20 | 50.8 (6.8)<sup>a</sup> | 95.7 (3.7)<sup>a</sup> | 66.7 (2.7)<sup>a</sup> | 94.3 (5.5)<sup>a</sup> | 92.7 (2)<sup>a</sup> | 80 (7.7)<sup>a</sup> |
| 25/15 | 69.2 (2.9)<sup>b</sup> | 100 (1.1)<sup>a</sup> | 86.7 (4.7)<sup>b</sup> | 100 (9.8)<sup>a</sup> | 100 (1.7)<sup>b</sup> | 76.7 (11.4)<sup>a</sup> |
| 17/7 | 60 (2.9)<sup>ab</sup> | 97.8 (2.7)<sup>a</sup> | 88.3 (4.2)<sup>b</sup> | 77.1 (9.8)<sup>a</sup> | 100 (1.7)<sup>b</sup> | 66.7 (5.4)<sup>a</sup> |
| *GA<sub>3</sub>* |  |  |  |  |  |  |
| 30/20 | 86.2 (6.6)<sup>**</sup> | 98.9 (1.8)<sup>n.s.</sup> | 63.3 (4.3)<sup>n.s.</sup> | 97.1 (7.4)<sup>n.s.</sup> | 96.9 (2)<sup>n.s.</sup> | 86.7 (8.6)<sup>n.s.</sup> |
| *After ripening* |  |  |  |  |  |  |
| 30/20 | 94.9 (4.9)<sup>**</sup> |  |  |  |  |  |
| 25/15 | 93.5 (6.2)<sup>*</sup> |  |  |  |  |  |
| 17/7 | 98.9 (3.6)<sup>***</sup> |  |  |  |  |  |
| *GA<sub>3</sub> + after ripening* |  |  |  |  |  |  |
| 30/20 | 93.5 (2.3)<sup>n.s.</sup> |  |  |  |  |  |

### Effect of temperature on seed germination rates

Non-dormant species germinated (T_min_) rapidly and within six days in at least one diurnal temperature treatment (Fig. 4). The two *Maireana* shrubs reached maximum VAG quickest, within one to five days, while tree species were slower to germinate. Time to maximum germination (T_max_) was significantly affected by temperature for all species except for *C. pauper*, with a general trend of decreasing germination times as diurnal temperatures increased (Table 3). T_max_ was significantly longer in *M. pyramidata* and *A. oleifolius* at 17/7°C and 25/15°C than at 30/20°C (P ≤ 0.014). The species with germination times least affected by temperatures were *H. tephrosperma* which had significantly delayed germination only at the coldest temperature (17/7°C; P = 0.014), and *C. pauper*, which had no significant change in germination times across all temperatures (P = 0.25).

**Table 3.** Time to minimum, 50% and maximum germination of seeds. incubated at 30/20°C, 25/15°C and 17/7°C (± standard error). *Atriplex rhagodioides* (AR) refer to seeds rendered non-dormant through a 12 month after-ripening.

| Species | Mean T<sub>min</sub> |  |  | Mean T<sub>50</sub> |  |  | Mean T<sub>max</sub> |  |  |
| --- | --- | --- | --- | --- | --- | --- | --- | --- | --- |
|  | 17/7°C | 25/15°C | 30/20°C | 17/7°C | 25/15°C | 30/20°C | 17/7°C | 25/15°C | 30/20°C |
| *Alectryon oleifolius* | 16.0 (0.9) | 6.5 (0.5) | 5.6 (0.5) | 20.3 (0.9) | 12.3 (2.7) | 8.8 (1.7) | 21.8 (1.0) | 14.8 (1.8) | 10.3 (1.3) |
| *Casuarina pauper* | 6.5 (0.3) | 3.3 (0.3) | 2.0 (0.0) | 9.0 (0.6) | 4.5 (1.2) | 3.3 (0.3) | 9.8 (0.6) | 7.5 (1.0) | 7.5 (1.04) |
| *Hakea tephrosperma* | 7.8 (0.3) | 4.8 (0.3) | 4.3 (0.5) | 10.0 (0.0) | 7.3 (0.3) | 6.8 (0.3) | 11.3 (0.3) | 9.3 (0.8) | 8.5 (0.5) |
| *Atriplex rhagodioides* | 14.0 (1.8) | 8.3 (0.9) | 7.8 (0.9) | 24.8 (1.0) | 18.5 (1.0) | 16.5 (1.8) | 26.0 (0.4) | 21.8 (0.3) | 20.0 (0.6) |
| *Atriplex rhagodioides* (AR) | 6.0 (0.0) | 3.0 (0.0) | 3.0 (0.0) | 8.7 (0.3) | 5.0 (0.0) | 4.7 (0.3) | 12.7 (0.3) | 9.7 (0.3) | 9.0 (0.0) |
| *Maireana pyramidata* | 1.5 (0.3) | 1.0 (0.0) | 1.0 (0.0) | 3.8 (0.3) | 2.5 (0.29) | 3.3 (0.3) | 5.0 (0.0) | 4.0 (0.0) | 3.5 (0.3) |
| *Maireana sedifolia* | 1.0 (0.0) | 1.0 (0.0) | 1.0 (0.0) | 2.3 (0.3) | 1.3 (0.3) | 1.3 (0.3) | 4.0 (0.0) | 2.8 (0.3) | 2.0 (0.0) |

**Fig. 4.**
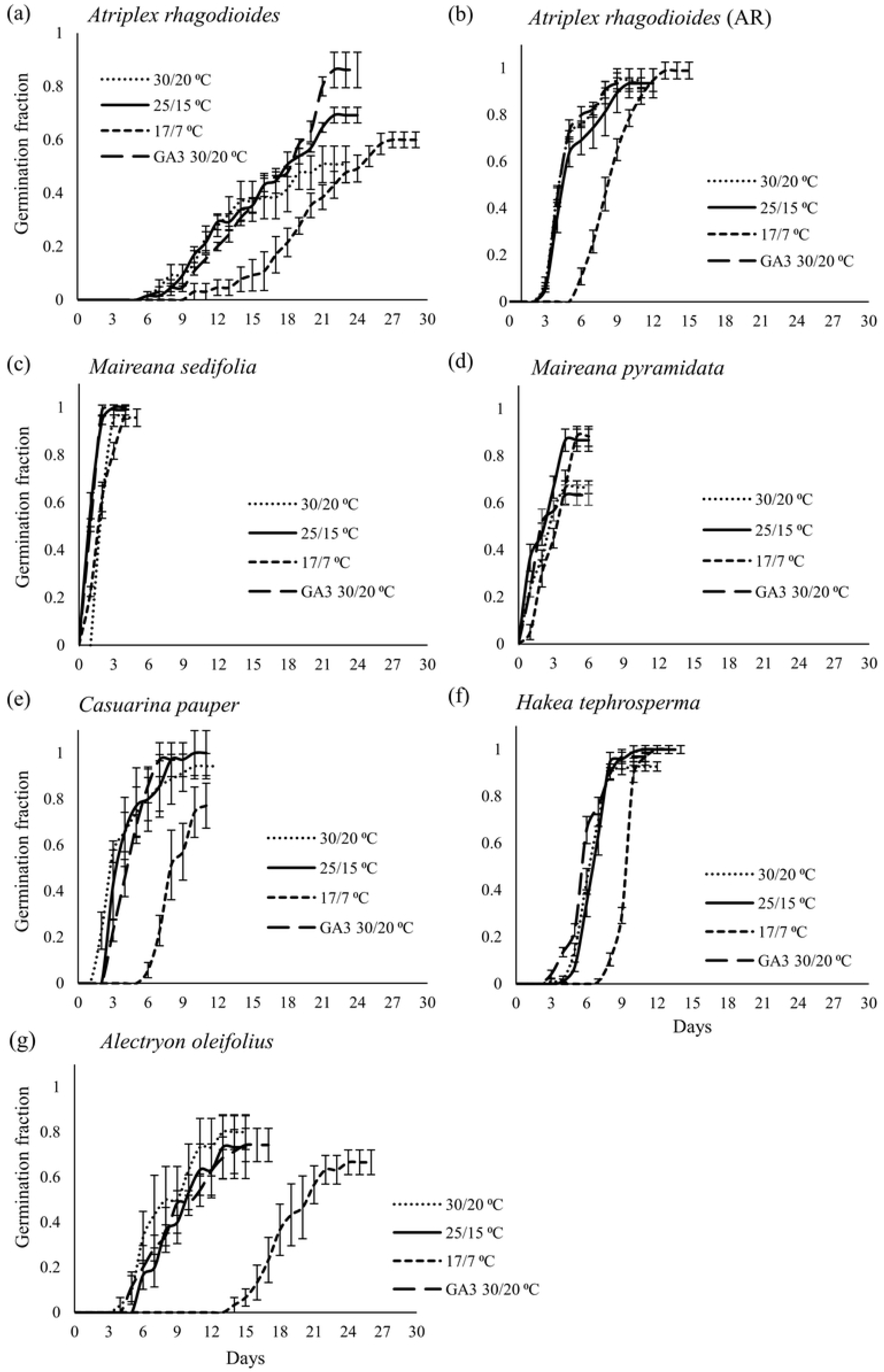
Cumulative germination (mean ± se) across diurnal temperatures: Seeds were incubated at 30/20°C, 25/15°C and 17/7°C for 30 days. Data also includes results from seeds incubated at 30/20°C, with the addition of a growth stimulant (GA_3_).

### Germination responses to GA_3_ and after-ripening

The only species with a positive germination response to GA_3_ treatments was *A. rhagodioides* (P = 0.010; P > 0.5 for all other species). *A. rhagodioides* achieved T_max_ in nine days with after-ripened seed compared to twenty days with fresh seed (Fig. 4b; Table 3). Similarly, for *A. rhagodioides* maximum VAG of after-ripened seed was significantly higher than for fresh seed (P ≤ 0.04 for all temperatures), and it eliminated the effect of GA_3_ treatments (P = 0.36) and widened temperature ranges for maximum germination, indicating dormancy loss.

### Seed longevity

*Myoporum platycarpum* was the only species to exhibit characteristics of recalcitrant seeds, meaning they do not survive desiccation or storage. Seeds of *M. platycarpum* were freshly collected and most had no embryo, and of the few that had an embryo most were non-viable. Seed freshly picked from adult trees showed only 13% viability, which fell to half that within one month of storage, and was close to 0% viability within six months of storage (Fig. 5). Most species showed a substantial decline in viability (>50%) within 12 months of storage, with the exception of *H. tephrosperma, A. rhagodioides* and *C. pauper*. These three species experienced less than 20% decline in viability within 12 months. The only species to show a decline in seed viability of <10% during 24 months in storage was *H. tephrosperma*. P50 for each species was (in order of longest to shortest lived, in months): *H. tephrosperma*, 84.1; *A. rhagodioides*, 32.0; *C. pauper*, 19.7; *M. sedifolia*, 14.7; *M. pyramidata*, 11.5; *G. parviflora*, 10.7; *A. oleifolius*, 8.7; *M. platycarpum*, 3.0.

**Fig. 5.**
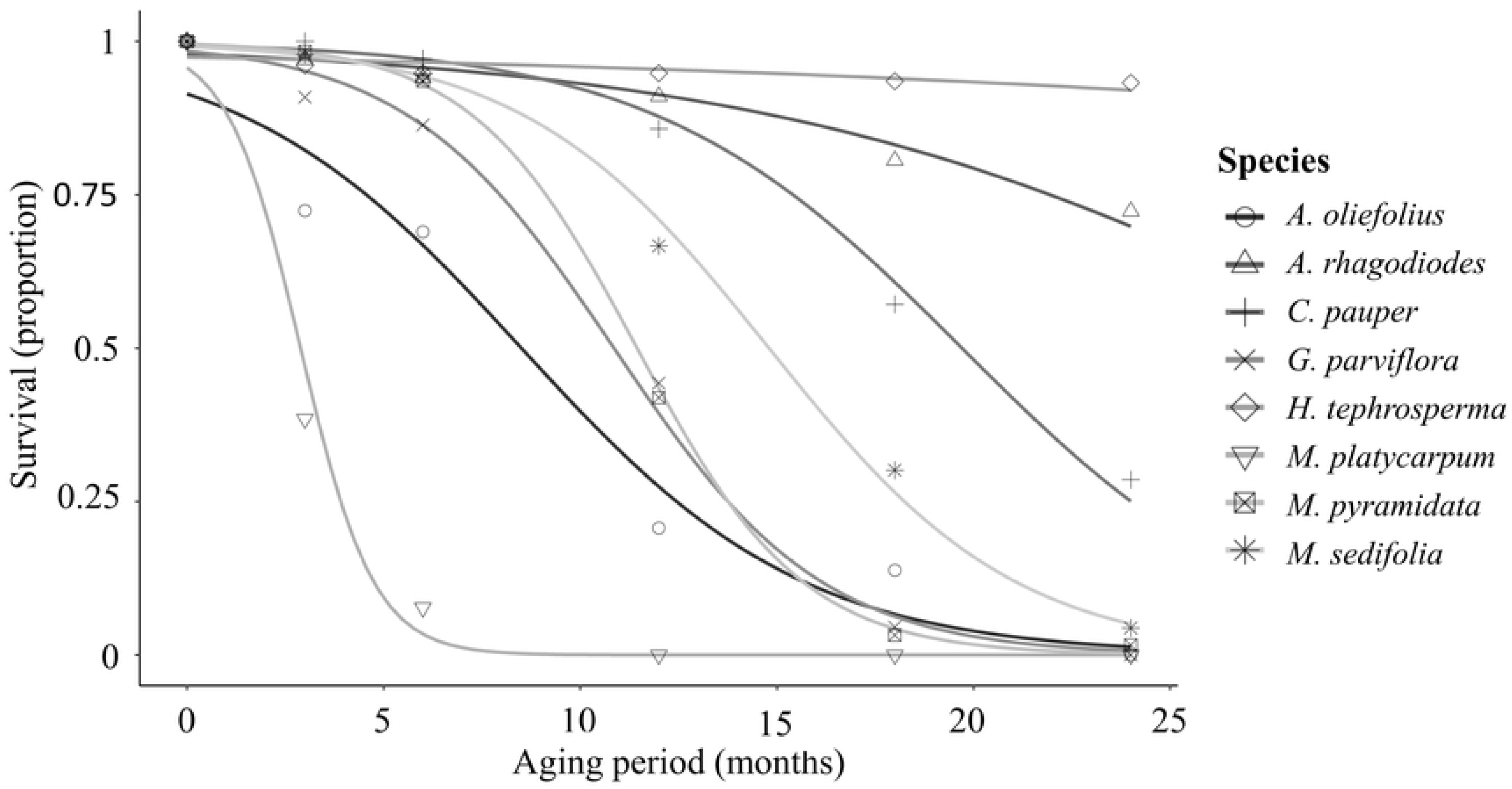
Loss of seed viability with time in storage. Seed age at beginning of experiment are show in Table 1.

## Discussion

Arid species in this study were generally categorised by two types of adaptive strategies to facilitate seed germination in sporadic rainfall: 1) rapid germination across wide diurnal temperatures, or 2) dormancy and/or long-lived seeds to temporarily delay, or stagger, germination. Our study demonstrates that rapid germination is a common, alternative and important strategy in seeds from arid zones, which allows seeds to capitalise on sporadic rainfall. Seeds of three species (*M. pyramidata, A. rhagodioides* and *H. tephrosperma*) had significantly higher VAG at cooler temperatures, than at 30/20°C, hence avoid germination when evaporation rates are highest [49]. Germination inhibition at hottest temperatures is considered an important germination strategy for seeds with dormancy or longevity in this study. We also demonstrated the tendency for seed weight and embryo type to be correlated to germination rate of the species in our study. Conversely, seed mass was unrelated to longevity and seed fill.

### Rapid germination across wide diurnal temperatures

All non-dormant seeds of species in this study exhibited rapid germination rates, which suggests that rapid germination is particularly well suited to this environment and may be at a selective advantage over other seed germination strategies. All species, except for dormant seeds of *G. parviflora* and *A. rhagodioides*, commenced germination within six days upon wetting and displayed T_50_ responses of ≤9 days at the highest temperatures tested. Rapid germinating species are able to take advantage of single rainfall events, whereas slow germinators require several rainfall events persisting across days [39], which rarely occurs in the arid zone studied here [49]. All species achieved >50% germination across all temperatures tested, suggesting they have wide thermal ranges for germination. Wide thermal ranges for germination allow seeds to take advantage of stochastic rainfall events that occur during any season. While this may enable opportunistic plant recruitment, additional studies should quantify hydrothermal niches for seed germination to further understand recruitment timing across different seasons.

Germination strategies, particularly those that affect the timing of germination such as dormancy, may be important determinants of population dynamics in arid ecosystems. For example, vegetation classification in this arid zone prior to disturbance was Belah-Rosewood Woodland and Belah-Bluebush Woodland [50], of which the dominant species include the non-dormant, fast germinators *C. pauper* and *Maireana* shrubs. The two species that have seeds with physiological dormancy, *A. rhagodioides* and *G. parviflora*, and non-dormant seeds of *A. oleifolius*, all of which had the slowest germination speeds, are less dominant [51] and appear as scattered individuals throughout the landscape [50]. Although the age of seeds in this study varied and pre-storage components may have contributed to aging, longevity or dormancy loss in some species, reports of non-dormancy here are consistent with the findings of Callister (2004) for all species, except *M. platycarpum*. Further studies are required to confirm dormancy in fresh seeds of *C. pauper, H. tephrosperma* and *A. oleifolius*. The lack of seeding events in these species during this study (potentially due to drought conditions) suggests that acquiring the quantities of seed required for their restoration may become more challenging under climate change.

Rapid germination appears to be unrelated to other morphological seed traits, except for seed mass and embryo type. All seeds in this study had fully developed embryos without an endosperm, or with embryos that are coiled and larger than the seed, and these are traits that are generally thought to indicate a rapid germination strategy [28, 29]. Germination for seeds with peripheral embryos (including *Atriplex* and *Maireana*) is typically very fast because it involves merely the uncoiling of the spiral embryo upon imbibition, which ruptures the seed coat [52]. Many such species with peripheral embryos inhabit high-stress environments, where the rapid exploitation of temporarily favourable conditions for germination is more important [40]. Our results showed that the size and development of embryos was not a consistent predictor of germination strategy.

The species in this study with the fastest germination rates (*C. pauper, M. sedifolia* and *M. pyramidata*), are dominant species from the region, and produced small seeds that were easier to obtain due to frequent and prolific seeding events. High seed production requires high maternal input but the risk of population crashes are mediated because these species are less dependent on high seed survival rates [1]. Other studies report strong evidence for survival advantages associated with larger seeds under stressful conditions [53-55] but, considering the variable size of arid seeds and the trade-off associated to increased seed production in small seeded species, survival advantages of large-seeded species does not appear large enough to counterbalance the advantage of small-seeded species during seed production [11, 56, 57]. We found that small seeded species were amongst the fastest to germinate, however the negative relationship between germination speed and seed size was weak, suggesting that seed size may not always be a reliable proxy for germination rate. Future studies are required to test a larger number of species of a greater order of magnitude of seed mass variation, prior to making general assumptions about the relationship between seed mass, germination rate and success.

### Seed dormancy and bet-hedging

Most arid seeds have dormancy traits to enable germination to coincide with periods of highest water availability [58, 59], yet only two species in this study (*A. rhagodioides* and *G. parviflora*) showed dormancy traits. These two species exhibited physiological dormancy traits as they imbibe water and have fully developed embryos, but failed to germinate to maximum potential within two weeks without treatment [59]. *A. rhagodioides* also showed a positive response to GA_3_. Fewer species showed signs of dormancy than was expected, though this is likely due to the unpredictability of rainfall at the study site. There is potential that due to the age of seeds used for this study, some of the study species may have exhibited dormancy traits as fresh seed. We have classified four such species (*A. oleifolius, C. pauper, H. tephrosperma* and *M. pyramidata*) as non-dormant, and we do not believe there would be significant levels of dormancy in the fresh seed of these species. Callister’s study [60] provides supporting evidence for this assertion for *C. pauper* and *A. oleifolius.* Local nurseries (including The Seeds of South Australia Database) and seed practitioners also support the claims of non-dormancy in these species (I. Sluiter, A. Quamby, T. Langdon, pers. comm. 2018). Traits that increase the likelihood of germination coinciding with periods of highest water availability may be less important in environments with unpredictable wetting pulses.

Delayed germination, through dormancy and/or bet-hedging, has been observed in many arid zone species [e.g.; 30, 31, 32]. Similarly, for *A. rhagodioides*, a small proportion of seeds germinate upon maturity, or upon seed dispersal, but germination rate and proportion improve with seed aging. This suggests that seeds can remain in the soil or canopy for years and stagger germination across seasons, which spreads the risk of recruitment failure through time and increases the probably that favourable conditions for seedling establishment will occur during a seeds’ lifespan [30, 61]. Physiological dormancy in *A. rhagodioides* was relieved through a period of after-ripening, also reported in other *Atriplex* species [62, 63]. After-ripening also enables a wider temporal window for germination which suggests that germination opportunities increase as seed ages. Developing a short-term soil seedbank, while waiting for suitable rainfall events, is an important adaptation in response to the unpredictability of resource availability in arid ecosystems. However, *A. rhagodioides* was the only dormant species studied, and non-dormant and prolifically-seeding trees are the most dominant species of the region. This again suggests that in arid zones without a wet season, playing it safe through dormancy and possibly bet-hedging may be less important than previously assumed.

### Seed longevity

Longevity is critical for species with dormancy traits or serotinous species with long-lived canopy-stored seed because they may need to survive multiple seasons. For example, some *Hakea* species develop a canopy-stored seedbank with fruit remaining closed for at least five years after maturity [64], while others are only weakly serotinous and release their seed cohort annually [65]. *Hakea tephrosperma* seeds in this study displayed exceptionally high longevity under laboratory conditions, despite possibly being non-dormant, suggesting seeds can persist is the canopy or soil for many years following seed maturity. Serotiny levels in *H. tephrosperma* are not yet reported, although only one occurrence of seed release was observed in populations from the study site during the two years of this study (personal observation). Our results are consistent with other studies that have found serotinous species typically have orthodox seeds that are long-lived [66], or desiccation tolerant.

Longevity results generated through controlled conditions may only reflect seed-storage behaviour during *ex situ* storage and may not represent results from soil-seed banking studies. Nevertheless, these results are directly applicable to the collection and storage of seed to be used in restoration efforts and, despite the limitations of the tests, provide insight into the longevity of these seeds that we might expect to observe in *in situ* conditions. Other studies report no soil seedbanks were maintained for *C. pauper* [67] and *M. sedifolia* [68]. *Ex situ* seed storage under dry conditions enhances seed longevity however, and species from arid environments are more likely to have comparatively long-lived seeds than those from cooler, wetter ecosystems [69]. Nonetheless, most species in this study had significantly higher longevity than expected [66, 69, 70]. High seed longevity is therefore likely to be a key adaptation to cope with the unpredictability of arid environments.

### Management implications

In arid zones, the requirement for wet conditions over long periods decreases the likelihood of seedling establishment. Tree species required wet conditions for longer than shrub species to germinate and *C. pauper*, the dominant tree species from the region, was the only tree species in this study to germinate within a few days, suggesting it to be a potential candidate for regeneration from seed-based restoration efforts. Restoration of slow-to-germinate species, such as *A. oleifolius* at winter temperatures, from seeding efforts alone may rely on engineering rehabilitation zones to increase temporal water availability. Seed viability and longevity was a critical problem for some species, in particular *M. platycarpum*, and restoration from seeding efforts may rely on further studies aimed at extending seed longevity through reduced temperature and humidity treatments.

While the germination responses of *M. platycarpum* were not classified in this study, it is reported to display T_50_ results within 4-5 days following wetting [60], consistent with other non-dormant trees species in this study. Callister, Florentine (71) suggests that seed-dormancy, not seed viability, is the key factor limiting germination in *M. platycarpum.* In contrast, as observed in our study, examining embryo firmness and colour through dissection microscopy is not an appropriate method to assess seed viability in *M. platycarpum* because TZ results show that healthy looking embryo are often dead (Figure 2f). Alternative viability testing, by assessing seed metabolic rate, may provide more accurate viability assessment [e.g.; 72] and should be considered for future testing. Our results suggest that seeds of *M. platycarpum* are non-dormant and that seed viability and short-lived seeds are the key factors inhibiting germination.

For dormant seeds, including *A. rhagodioides* and *G. parviflora*, dormancy should be alleviated prior to seeding to increase germination speed, provide wider temporal ranges for germination [73] and reduce the chances of infection and death. Due to lack of germination responses to treatments to relieve dormancy, and a lack of viable seed available for further germination treatments, dormancy cues of *G. parviflora* remain unresolved. Most species from the same family as *G. parviflora*, the Rutaceae family, produce seeds with physiological dormancy [59, 74-76]. Restoration of species with elusive dormancy cues may rely on tubestock based technologies but, for *G. parviflora*, we consider seed viability as a key factor limiting recruitment from seed. Approximately 90% of seeds fail to pass the seedling emergence stage [77] and meeting the demand for seed used in arid restoration will rely on better understanding the dormancy cues and germination niche of species [78-80].

## Conclusion

We discuss seed morphology and physiology, and the germination behaviour of seeds that may facilitate seed survival and growth in arid zones. Most species exhibit rapid germination across wide diurnal temperatures, often producing prolific quantities of seed, which enables species to take advantage of rainfall events that fall across all seasons. Fewer species tend to avoid unfavourable conditions by delaying germination through seed dormancy and possibly bet-hedging. High seed longevity under *ex situ* storage conditions was observed in most species included in this study, which has important implications for restoration. While seed dormancy is an important survival strategy for an arid seed, rapid germination and high seed production rates may be alternative regeneration traits and have a crucial role in explaining population dynamics in arid ecosystems with unpredictable rainfall. In such systems, the benefits of rapid germination may outweigh those associated with delayed germination through dormancy, and the population structure of remnant ecosystems in the region (which are dominated by rapid germinators) may be a testament to this.

## Acknowledgements

We wish to acknowledge the environmental team at Cristal Mining for instrumental support and access to land and to Kings Park Science for trainings in seed-based technologies. Finally we thank Dr Ian Sluiter and Tim Zweirsen for sharing insights to seed-based restoration approaches in arid zones and support for the project.

